# Deep-Interact Studio: An Interactive Deep Learning Model Building Platform for Biomolecular Interaction Prediction

**DOI:** 10.64898/2026.07.02.736034

**Authors:** Dipayan Sarkar, Koushik Bardhan, Chiranjib Sarkar

**Affiliations:** Computational Systems Biology Laboratory, Department of Bioinformatics, University of North Bengal, India

**Keywords:** biomolecular interaction prediction, customizable deep-learning models, multi-model comparative inference, model interpretability, web-based bioinformatics platform

## Abstract

**Motivation:** Deep learning has rapidly become essential for predicting biomolecular interactions; however, most web-tools expose only a single, pre-built model with a fixed, non-configurable architecture that users cannot redesign, retrain on their own data, or compare; they are typically dedicated to one interaction type and often one species, and report prediction scores with little interpretability. These constraints force researchers across several disconnected, single-purpose tools and limit the flexibility, reproducibility, and long-term usability of existing platforms.

**Results:** We present Deep-Interact Studio, a unified, web-based deep-learning platform that addresses these limitations by shifting interaction prediction from a model-centric to a user-driven, comparative, and interpretable paradigm. Within a single interface spanning all four interaction classes, namely protein-protein, drugtarget, RNA-protein, and protein-DNA, users design their own model architectures layer by layer, configure training hyperparameters, and train them on their own data, including custom, species-specific datasets. Multiple user-built models can then be trained under identical conditions and compared side by side at both the training and inference levels, while integrated interpretability, including SHAP-based feature attribution, embedding-space visualization, and interaction hub analysis, turns predictions into auditable, mechanistically grounded results. Deep-Interact Studio is, to our knowledge, the only such platform to combine fine-grained per-layer model customization with multi-model comparison and interpretability, offering a flexible and transparent alternative to fixed, single-purpose tools.

**Availability and implementation:** Deep-Interact Studio is freely available as a web application at https://deepinteract.compbiosysnbu.in/, with no login or installation required.

## 1 Introduction

Web-based computational tools have become indispensable in modern molecular biology, democratizing access to powerful predictive models for experimental researchers who lack specialized programming or high-performance computing resources [1]. Among the problems where such tools are most needed is the prediction of biomolecular interactions, which form the mechanistic basis of nearly all cellular processes from signaling and gene regulation to metabolism and immunity and whose dysregulation underlies diseases ranging from cancer to neurodegeneration [2–6]. Reliable prediction of the major interaction classes; protein-protein, drug-target, RNA-protein, and protein-DNA; is therefore a central goal in bioinformatics, with direct relevance to drug discovery and personalized medicine [7]. Because experimental screening of these interactions remains costly and time-consuming, there is a strong and growing need for accessible, accurate, and interpretable web-based prediction tools that bring these capabilities to experimental biologists without specialized computational expertise.

Deep learning has rapidly transformed this landscape [8–10]. Large interaction databases and modern neural architectures, especially pre-trained language models such as ESM-[11] for proteins and ChemBERTa [12] for small molecules [13, 14], have driven a proliferation of dedicated prediction tools. For protein-protein interactions, servers such as ProteinPrompt [15] and deep pipelines like DPPI [16] predict binding from sequence alone, while for drug-target interactions GraphDTA [17], DeepPurpose [18], Drug-Online [19], and DeepMolecules [20] predict binding and affinity. For RNA-protein interactions, PrismNet [21] and RBPsuite [22] predict binding from sequence and structure. More general sequence-analysis platforms; iLearnPlus [23], DeepBIO [24], and BioSeq-BLM [25]; together with RNA-focused tools such as EnrichRBP [26] and language-model servers such as ProteinBERT [27], expose transfer learning through fixed, pre-trained architectures.

Despite their utility, these tools share several recurring limitations. Most expose only a single pre-built model with a fixed, non-configurable architecture; the user can submit sequences and receive a prediction, but cannot alter the network design, substitute a different architecture, or build and compare alternative models on their own data. Compounding this, such models are frequently trained on a single benchmark dataset or a single species, so their reported performance does not necessarily transfer to new organisms, assay conditions, or distributions, leaving users with no way to retrain or adapt the model to their specific problem. These tools are also typically single-task, dedicated to one interaction type, so studying protein–protein, drug-target, and RNA-protein interactions forces the user to move across several disconnected tools, each with its own input format, interface, and output conventions. A further practical concern is longevity; a large fraction of previously published interaction web servers are no longer accessible online, and many that remain offer limited interpretability, reporting a bare prediction score without feature attribution [28] or embedding-space context [29]. Together, these limitations restrict the flexibility, reproducibility, and long-term usability of existing interaction-prediction platforms.

To address these needs, we present Deep-Interact Studio, a unified, web-based deeplearning platform for biomolecular interaction prediction that shifts the paradigm from model-centric to comparative, data-centric and user-driven analysis. Within a single interface covering protein-protein, drug-target, RNA-protein, and protein-DNA interactions, users are no longer limited to a fixed, pre-built predictor but can design their own models and train them on their own data, including custom, species specific datasets, which is particularly valuable for the many lesser studied or non-model organisms that existing single species tools do not serve well. Rather than committing to one architecture, users can create several named models, train them under identical conditions, and directly compare their predictive behaviour, examining and contrasting the inference performance of each model to identify the one best suited to their problem. Each model is further accompanied by both training and inference level insight and interpretability, so that predictions are not delivered as opaque scores but as auditable, mechanistically grounded results. By bringing custom model construction, multi-model comparison, cross-modality coverage, and integrated interpretability together in one accessible platform, Deep-Interact Studio aims to transform the way researchers approach biomolecular interaction prediction.

## 2 Materials and Methods

Deep-Interact Studio is a fully web-based, no-code platform built on a layered, service-oriented architecture [30]. A Streamlit interface [1] communicates with a FastAPI backend that handles validation and job management, while long-running embedding and training tasks are dispatched to GPU-accelerated Celery workers through a Redis broker and persisted, together with job metadata, in PostgreSQL. Each submission is registered under a unique run identifier and processed asynchronously, allowing users to track status and retrieve results without blocking the interface. The full stack is containerized with Docker Compose and served behind nginx; deployment details and server specifications are given in Supplementary Section S1.

The platform implements a modular, end-to-end pipeline organized into five stages; data preparation, model training, model insights, inference, and visualization (Figure 1, supplementary Figure S1). Uploaded sequence and chemical data are validated, cleaned, and split into stratified training and test partitions, with automatic generation of balanced negative pairs when datasets are class-imbalanced. Inputs are then encoded into fixed representations using frozen pre-trained foundation models; ESM-2 [11] for proteins and ChemBERTa [12] for compounds, with an RNA-specific encoder for RNA-protein tasks. From these embeddings, users construct one or more classifiers through an interactive, layer-by-layer model builder that supports linear, convolutional (CNN1D), recurrent (BiLSTM, GRU), transformer, and residual blocks, each with per-layer control of dimensionality, activation, dropout, and normalization, alongside a real-time architecture preview and parameter-count estimate (Supplementary Table s1). Multiple models are trained simultaneously under identical data partitions and hyperparameters, and every run is recorded in a model registry that stores the architecture, configuration, metrics, and learned embeddings to guarantee full reproducibility (detailed in supplementary s2).

**Figure 1:**
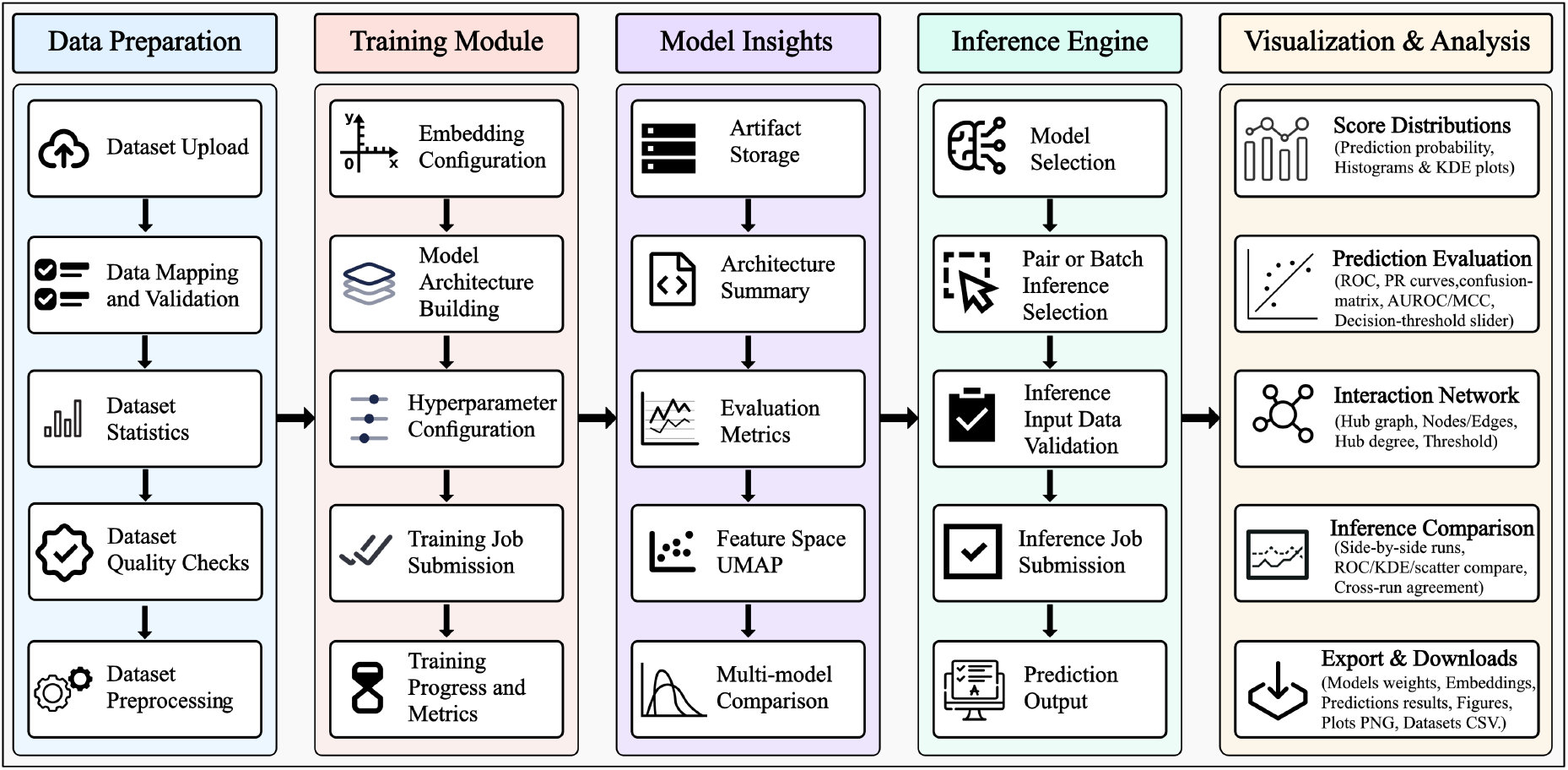
Overview of the Deep-Interact Studio workflow for unified multi-model training, comparative inference, and interpretability.

Trained models can be applied through single- or multi-model inference, in which up to five models predict on a common input and their outputs are aligned at the sample level for direct comparison. Beyond aggregate performance metrics, the platform quantifies per-sample agreement and ranks the drug-target pairs on which models most strongly disagree, directly flagging uncertain predictions for experimental follow-up. Model behavior is further interpreted through SHAP-based feature attribution [28], UMAP projection of the input and learned feature spaces [29], and an interaction-network view that highlights high-degree hub targets. All quantitative results, plots, and ranked tables are exportable in machine-readable formats.

The platform was benchmarked on curated PPI, RNA-protein, and drug-target interaction datasets, and model performance was assessed using accuracy, precision, recall, specificity, F1-score, Matthews correlation coefficient, AUROC, and AUPRC. Full dataset composition and metric definitions are provided in Supplementary Sections s1.4 and s1.5.

## 3 Results

We validated Deep-Interact Studio through an end-to-end drug–target protein interaction (DTPI) case study performed entirely through the web interface without any scripting; the full workflow, figures, and metric tables are provided in Supplementary Section S2. Three architectures sharing frozen ESM-2 and ChemBERTa encoders and a common input projection, but differing in a single hidden layer (linear, BiLSTM, and residual), were registered and trained in parallel over 30 epochs and converged stably to comparable discriminative performance (AUROC 0.880–0.889, AUPRC 0.892–0.897; Supplementary Figure S3, Tables S3–S4). The multi-model inference engine then executed three independent inference runs on a shared test set, automatically ranking them across accuracy, AUROC, AUPRC, F1, and MCC and flagging the best run (Supplementary Figure S5, Table S5). Beyond aggregate metrics, the platform’s per-pair disagreement score (the standard deviation of predicted probability across runs) identified high-uncertainty drug–target pairs for experimental follow-up (Supplementary Figure S7, Table S6), while SHAP attribution, UMAP projections of the input and learned feature spaces, and a degree-ranked interaction hub network exposed the representational basis of the predictions (Supplementary Figure S8). All metrics, plots, and ranked tables were exported in machine-readable form directly from the interface. Relative to existing sequence-based interaction and model-building platforms, Deep-Interact Studio is, to our knowledge, the only tool that combines no-code multi-architecture building, side-by-side multi-model training and inference comparison, interpretability, and reproducibility across all four interaction types in a single web application (Table 1); an expanded, category-wise qualitative comparison against these platforms is provided in Supplementary Table S8.

**Table 1:** Feature comparison of Deep-Interact Studio (DIS) with existing platforms: DeepPurpose (DP) [18], iLearnPlus (iLP) [23], DeepBIO (DB) [24], BioSeq-BLM (BSL) [25], Drug-Online (DO) [19], DeepMolecules (DM) [20]. ✓ = supported, ~ = partial, × = not supported.

| Feature / Capability | DP | iLP | DB | BSL | DO | DM | DIS |
| --- | --- | --- | --- | --- | --- | --- | --- |
| Web-based, no-code interface | × | ✓ | ✓ | ✓ | ✓ | ✓ | ✓ |
| GPU-accelerated training | ✓ | × | ✓ | × | ~ | × | ✓ |
| All four interaction types (PPI/DTPI/RPI/PDI) | × | × | × | × | × | × | ✓ |
| User-defined model architecture | × | ~ | ✓ | ~ | × | × | ✓ |
| Per-layer activation/dropout/norm control | × | × | × | × | × | × | ✓ |
| Live training metrics / loss curves | ✓ | ✓ | ✓ | × | × | × | ✓ |
| Multi-model training comparison (up to 5) | × | ✓ | ✓ | ✓ | × | × | ✓ |
| Multi-model inference comparison (up to 5) | × | × | × | × | × | × | ✓ |
| Interpretability (UMAP / SHAP) | × | × | ~ | × | × | × | ✓ |
| Reproducible registry / job tracking | × | ~ | ~ | × | ✓ | × | ✓ |

## 4 Conclusion

Deep-Interact Studio reframes biomolecular interaction prediction as a customizable, comparative, and interpretable task rather than a fixed, single-model one. Within one web-based interface, users build their own classifier architectures layer by layer, configure training hyperparameters, and train models on their own data, including custom, species-specific datasets, across four interaction classes, namely protein-protein, drugtarget, RNA-protein, and protein-DNA. As demonstrated in our case study, several user-built models can be trained under identical conditions and compared side by side at both the training and inference levels, with per-sample disagreement analysis and integrated interpretability, including SHAP-based feature attribution, embedding-space visualization, and interaction hub analysis, turning predictions into auditable, mechanistically grounded results that are fully reproducible through a central model registry.

Because the platform is built around a modular, customizable paradigm, it is readily extensible. Future development will broaden the model builder with additional deeplearning layer types, expanded hyperparameter and training-strategy options, and a wider selection of pre-trained embedding models, as well as support for further interaction types. More broadly, the same customizable, comparative, and interpretable framework can be stretched beyond interaction prediction to other sequence- and structure-based biological prediction tasks, providing a general foundation for transparent and reproducible deeplearning analysis in bioinformatics.

## Author contributions

Dipayan Sarkar (Conceptualization, Methodology, Software, Validation, Formal analysis, Investigation, Data curation, Visualization, Writing—original draft, Writing review & editing), Koushik Bardhan (Methodology, Validation, Investigation, Writing review & editing), and Chiranjib Sarkar (Conceptualization, Supervision, Project administration, Resources, Writing review & editing).

## Supplementary Data

Supplementary information is provided in the accompanying Supplementary File.

## Data availability

The datasets used in this study are freely available at https://deepinteract.compbiosysnbu.in/benchmark_datasets. The project is open source and available on GitHub at https://github.com/Dipayan26/Deep-interact-Studio.

