## Supplementary Information for "Deep-Interact Studio: An Interactive Deep Learning Model Building Platform for Biomolecular Interaction Prediction"

### S1 Materials and Methods

#### S1.1 System Architecture

The architecture summarized in the main text is organized into four cooperating layers [1] (Figure S2), a presentation layer [2], an application layer that additionally enforces rate limiting, a GPU-accelerated worker layer, and a database and infrastructure layer that pairs PostgreSQL job metadata with file-volume storage for models, embeddings, and results. Job handling is fully asynchronous; on submission the backend registers the job, returns immediately, and the worker updates its lifecycle state (queued/running/completed/failed) and outputs against the run identifier, which the frontend polls for status and results. Running each layer as an isolated container cleanly separates user interaction, API logic, background computation, and persistence, which is what allows the platform to scale across concurrent submissions while remaining maintainable.

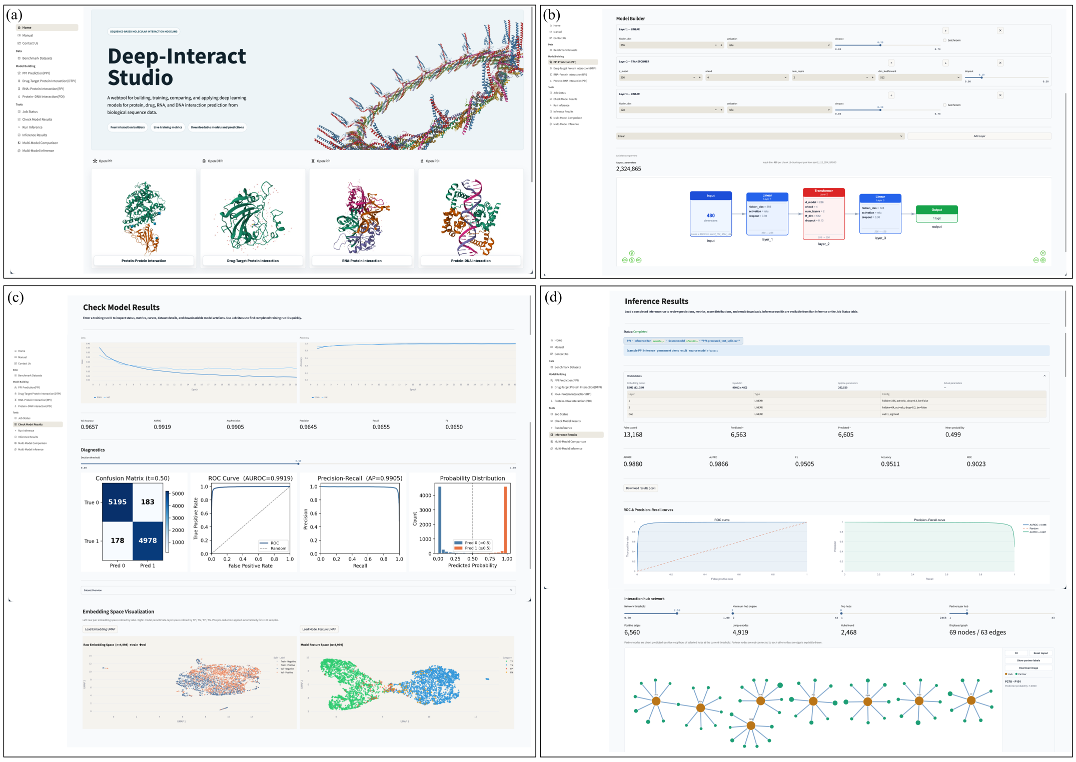

Figure S1: Overview of the Deep-Interact Studio web interface. (a) Home page, providing entry points to the four sequence-based interaction builders. (b) No-code model builder, in which classifier layers (e.g. linear, transformer) are stacked and configured interactively, with a live architecture preview and real-time parameter-count estimate. (c) Model results page, reporting training/validation loss and accuracy curves, summary metrics. (d) Inference results page, showing per-run prediction summaries and metrics, downloadable predictions, and an interaction hub network

#### S1.2 Server description

Deep-Interact Studio is hosted by the Computational Systems Biology Lab, Department of Bioinformatics, University of North Bengal, and is freely available at `https://deepinteract.compbiosysnbu.in/`. The application runs on an Ubuntu 24.04 LTS server (Intel i5-12400F, 32 GB RAM) served behind nginx 1.24.0, with the full stack (Streamlit, FastAPI, Celery, Redis, and PostgreSQL) orchestrated through Docker Compose for asynchronous job submission, execution, and result management. The interface was tested on Google Chrome, Microsoft Edge, and Mozilla Firefox.

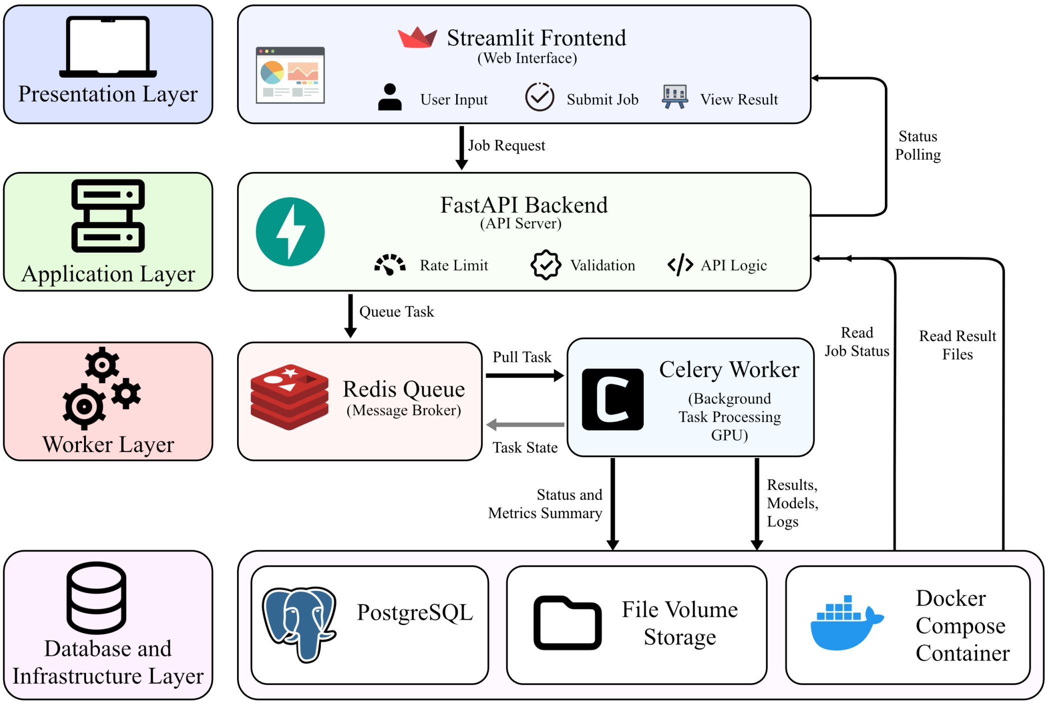

Figure S2: Schematic representation of the end-to-end system architecture and job processing pipeline of Deep-Interact Studio.

#### S1.3 Workflow

The five-stage pipeline outlined in the main text is described in detail below, one subsection per stage, from data ingestion through to interpretability and reporting.

##### S1.3.1 Data acquisition and preprocessing

The workflow begins with the ingestion of input data, comprising biological sequences and chemical structural information. A standardized preprocessing pipeline is applied to ensure consistency across all downstream analyses. This includes data cleaning and sampling, stratified train-test splitting, and selection of positive and negative pairs over the whole population. In parallel, dataset-level statistics are computed to characterize the input data distribution, including class imbalance, feature summaries, missing value patterns, and correlation structures, to provide an initial understanding of the dataset and inform subsequent modeling decisions. When the data lack negative pairs or are class-imbalanced, the system generates negative samples through randomized pairing of non-interacting entities to obtain a balanced dataset.

##### S1.3.2 Foundation-model embedding

Validated inputs are encoded into fixed-length representations using frozen, pre-trained foundation models: ESM-2 [3] for protein sequences and ChemBERTa [4] for compound SMILES, with an RNA-specific encoder for RNA-protein tasks. Embeddings can be mean-pooled into a single vector or retained as windowed sequence tensors, and interacting partners are combined by concatenation, element-wise product, absolute difference, or a combination thereof before being passed to the trainable classifier head. Holding the encoders frozen keeps the downstream parameter budget small and ensures that comparisons across architectures reflect differences in the classifier rather than in the representation.

##### S1.3.3 Multi-model training and model registry

Following preprocessing, multiple deep learning models can be trained on the same dataset under identical experimental conditions. The platform supports diverse architectures (Table S1), including linear models, 1D convolutional neural networks (CNN1D), recurrent neural networks (e.g. BiLSTM, GRU), transformer-based models, and residual architectures. A centralized training orchestrator manages hyperparameter configurations, training schedules, and resource allocation to ensure controlled and comparable experiments. Users can build the model layer-by-layer and customize each layer based on their requirements, previewing the model architecture and parameter count in real time. Each trained model is subsequently stored in a model registry, which maintains comprehensive metadata, including model architecture, dataset identifiers, training configurations, performance metrics, predicted outputs, learned embeddings, and runtime statistics, enabling versioning, traceability, and reproducibility of experiments.

##### S1.3.4 Multi-model inference and comparative analysis

The inference module supports both conventional single-model prediction and a multi-model inference mode, in which multiple trained models are applied to the same input dataset. Predictions are aligned at the sample level to enable direct comparison, and a comparison engine aggregates outputs from all selected models and evaluates them using standard performance metrics such as accuracy, sensitivity, specificity, and area under the curve (AUC). Beyond aggregate metrics, the platform conducts prediction agreement analysis, identifying regions of consensus and disagreement among models, ranking difficult samples by their per-pair disagreement score (the standard deviation of predicted probabilities across runs), and comparing per-model confidence scores to assess prediction reliability.

###### S1.3.5 Interpretability and data-centric analysis

To facilitate deeper understanding of model behavior, the workflow incorporates a data-centric interpretability layer. Dataset-level insights are revisited alongside model outputs to contextualize performance outcomes. Feature distributions are analyzed to assess class separability, while dimensionality reduction via Uniform Manifold Approximation and Projection (UMAP) is applied to both the raw input embeddings and the learned feature space to visualize clustering patterns, overlap regions, and latent structure. SHAP-based feature attribution quantifies the contribution of individual embedding dimensions and encoder subspaces, and a network visualization module renders predicted interactions as a hub network with degree-centrality node sizing.

###### S1.3.6 Output generation, reporting, and reproducibility

An interactive dashboard allows users to visualize comparative performance metrics, inspect inference-level predictions, and analyze feature-space representations. Results can be exported as tabular data (CSV/Excel), plots, and reproducible, publication-ready summaries that encapsulate model configurations, evaluation metrics, and analytical findings. Throughout the workflow, reproducibility is maintained via consistent preprocessing, controlled experimental settings, fixed random seeds, and comprehensive logging keyed to each run identifier. The modular architecture allows seamless integration of new models and datasets, enabling a transition from isolated model development to systematic, comparative, and interpretable multi-model analysis.

Table S1: Configurable layer blocks available in the Deep-Interact Studio no-code model builder. Any blocks may be stacked layer-by-layer into a custom classifier head, followed by a final linear layer that outputs the interaction probability.

| **Layer Block** | **Primary Functional Role** | **Key Configurable Parameters** |
|---|---|---|
| Linear | Baseline classification and feature integration | Hidden dimension, activation, dropout, batch normalization |
| CNN1D | Local motif and short-range sequence pattern extraction | Output channels, kernel size, activation, dropout |
| BiLSTM | Bidirectional sequential dependency learning | Hidden size, number of layers, dropout |
| GRU | Gated sequential dependency learning | Hidden size, number of layers, bidirectionality, dropout |
| Transformer | Context-aware global modeling via self-attention | Model dimension, attention heads, number of layers, feed-forward dimension, dropout |
| Residual | Stable deep feature propagation (skip connection + LayerNorm) | Hidden dimension, activation, dropout, batch normalization |

#### S1.4 Datasets

The platform was benchmarked on three curated collections of experimentally verified biomolecular interactions, one per task: protein-protein interaction (PPI) [5], RNA-protein interaction (RPI) [6], and drug-target protein interaction (DTPI) [7]. The PPI set comprises human protein pairs derived from primary amino acid sequences; the RPI set is drawn from NPInter across six organisms; and the DTPI set covers human drug-target pairs. In each case, positive interactions are balanced with constructed negative pairs. The exact composition, feature representation, and organism coverage of each dataset are given in Table S2.

Table S2: Comprehensive Summary of Dataset Composition, Dimensionality, Feature Representation, and Organism Coverage for PPI, RPI, and DTPI modules of Deep-Interact Studio.

| **Dataset** | **Data Dimensions** | **Key Features** | **Organism** |
|---|---|---|---|
| PPI | 32224 positive, Total 65836 | Conjoint triad (21-residue features); latent topic vectors (50–100 dims) | Human |
| RPI | 10412 positive, Total 20824 | RNA: $k$-mers, Zernike moments; Protein: PSSM ($20 \times L$ matrices) | Multi-organism (6 species) |
| DTPI | 13830 positive, Total 27464 | Drug: 2D fingerprints ($\sim$1,024 bits); Target: sequence homology graphs | Human |

#### S1.5 Evaluation Metrics

Model performance was comprehensively evaluated using an ensemble of standard binary classification metrics tailored for biomolecular interaction prediction tasks, including accuracy, precision, recall (sensitivity), specificity, F1-score, Matthew's correlation coefficient (MCC), area under the receiver operating characteristic curve (AUROC), and area under the precision-recall curve (AUPRC). These metrics provide multifaceted assessment of predictive capability, and are formulated:

$$\text{Accuracy (ACC)} = \frac{TP + TN}{TP + TN + FP + FN} \tag{1}$$

$$\text{Positive predictive value (Precision or PPV)} = \frac{TP}{TP + FP} \tag{2}$$

$$\text{Sensitivity (SN or recall)} = \frac{TP}{TP + FN} \tag{3}$$

$$\text{Specificity (SP)} = \frac{TN}{TN + FP} \tag{4}$$

$$F_1 = \frac{2 * Precision * Recall}{Precision + Recall} \tag{5}$$

$$MCC = \frac{TP \times TN - FP \times FN}{\sqrt{(TP + FP)(TP + FN)(TN + FP)(TN + FN)}} \tag{6}$$

$$\text{AUROC} = \int_0^1 \text{TPR } d(\text{FPR}) \tag{7}$$

$$\text{AUPRC} = \int_0^1 \text{Precision } d(\text{Recall}) \tag{8}$$

where $TP$, $TN$, $FP$, and $FN$ represent true positives, true negatives, false positives, and false negatives, respectively.

### S2 Results

This section provides the complete results of the drug–target protein interaction (DTPI) case study summarized in the Results section of the main text, including the full set of performance tables, comparison figures, and interpretability analyses generated through the Deep-Interact Studio web interface.

#### S2.1 Multi-Model Training and Comparative Inference

To demonstrate the usability and analytical functionality of the Deep-Interact Studio web platform, we applied the Drug-Target Protein Interaction (DTPI) prediction module to a binary interaction classification task. The workflow encompasses two sequential stages accessible through the platform's graphical interface: (i) multi-model parallel training with real-time performance visualization, and (ii) multi-run comparative inference with per-sample disagreement analysis. All experiments shared an identical module-wise input representation of dimension, constructed by concatenating chemical embeddings from ChemBERTa-zinc-base-v1 or RNA embeddings RNA-FM with protein sequence embeddings from ESM-2, all pre-trained encoders held frozen throughout training.

##### S2.1.1 Model Architectures and Configuration

Three distinct neural network architectures were registered and trained concurrently through the platform's Model Builder interface. All three models share an identical input projection layer (Linear, 256 hidden units, ReLU activation, dropout rate 0.3) and a sigmoid-activated binary output head, but differ exclusively in their second hidden layer, enabling a controlled architectural comparison under strictly identical data preprocessing and training conditions. The architectures are summarized in Table S3.

Table S3: Architectural overview of the three DTPI models trained via the Deep-Interact Studio multi-model interface.

| **Property** | **M1 (7111e32f)** | **M2 (aca39852)** | **M3 (efc533b6)** |
|---|---|---|---|
| Task | DTPI | DTPI | DTPI |
| Input Dim. | 1,248 | 1,248 | 1,248 |
| ~Parameters | 336,257 | 715,265 | 452,097 |
| Chemical Encoder | ChemBERTa-zinc-base-v1 | ChemBERTa-zinc-base-v1 | ChemBERTa-zinc-base-v1 |
| Protein Encoder | ESM-2 (35M) | ESM-2 (35M) | ESM-2 (35M) |
| Layer 1 | Linear(256, ReLU, d=0.3) | Linear(256, ReLU, d=0.3) | Linear(256, ReLU, d=0.3) |
| Layer 2 | Linear(64, ReLU, d=0.2) | BiLSTM(128, layers=1, d=0.3) | Residual(256, ReLU, d=0.3) |
| Output | Linear(1, Sigmoid) | Linear(1, Sigmoid) | Linear(1, Sigmoid) |

###### S2.1.2 Multi-Model Training Dynamics

All three models were trained simultaneously via the platform's Training Orchestrator over 30 epochs on the same preprocessed dataset partition. The web interface rendered real-time training curves for each registered model, enabling side-by-side monitoring of learning dynamics without requiring manual experiment management. Figure S3a presents the validation loss and validation accuracy trajectories across epochs for all three models and all three models exhibit consistent monotonic loss reduction and accuracy improvement, converging to comparable final performance with no evidence of overfitting. Both plots in Figure S3b reveal near-identical discriminative capacity across architectures, with AUROC values ranging from 0.8804 to 0.8893. The high average precision scores (0.8924-0.8969) confirm strong performance under class imbalance conditions. Figure S3c presents confusion matrices, which corroborate the metric-level findings. The evaluation metrics obtained on the validation partition. M2 achieves the highest true positive count (1,820) and lowest false negative rate, while M1 records less false positives i.e. 379. Model M1 (7111e32f) employs a conventional two-layer fully connected architecture (336,257 parameters) representing a compact linear baseline. Model M2 (aca39852) incorporates a bidirectional LSTM layer in place of the second linear layer, substantially increasing the parameter count to 715,265 and introducing recurrent sequence modelling capacity. Model M3 (efc533b6) adopts a residual connection block (256 units, 452,097 parameters), enabling gradient flow through the network whilst preserving the original feature representation. This design allows the platform to benchmark representational capacity against architectural depth on a common dataset. M2 achieved the highest true positive count (1,820), correctly classifying the greatest proportion of true interaction pairs, with a correspondingly low false negative count of 393. M1 exhibited the fewest false positives (379) consistent with its superior precision, while M3 produced intermediate results (1,773 true negatives, 1,742 true positives). The platform renders these matrices interactively, allowing users to inspect per-class error distributions for each registered model without additional scripting. All three architectures converged smoothly, with validation loss decreasing from approximately 0.58 at epoch 1 to approximately 0.43 by epoch 30, and validation accuracy improving from near 0.67 to approximately 0.80-0.81 across the same interval. M2 and M1 demonstrated marginally faster initial convergence relative to M3, though all three models reached comparable terminal accuracy. The absence of divergence or catastrophic loss spikes across any of the three architectures validates the stability of the platform's shared training pipeline and data loading procedures.

###### S2.1.3 Comparative Performance Metrics Following Training

Upon completion of training, the platform's Evaluation Engine automatically computed a comprehensive suite of classification metrics for each model on the held-out validation set. The results are presented in Table S4. All metric values were exported directly from the platform's tabular comparison interface. M2 achieved the highest AUROC (0.8893), AUPRC (0.8969), recall (0.8224), and F1 score (0.8121), suggesting that the bidirectional LSTM layer confers an advantage in capturing sequential interaction patterns within the concatenated embedding space. M1 attained the highest raw validation accuracy (0.8100) and precision (0.8226), reflecting its more conservative prediction behavior and lower false-positive rate. M3 produced uniformly competitive performance across all metrics, with the highest F1 and AUROC values exceeding those of M1, despite being

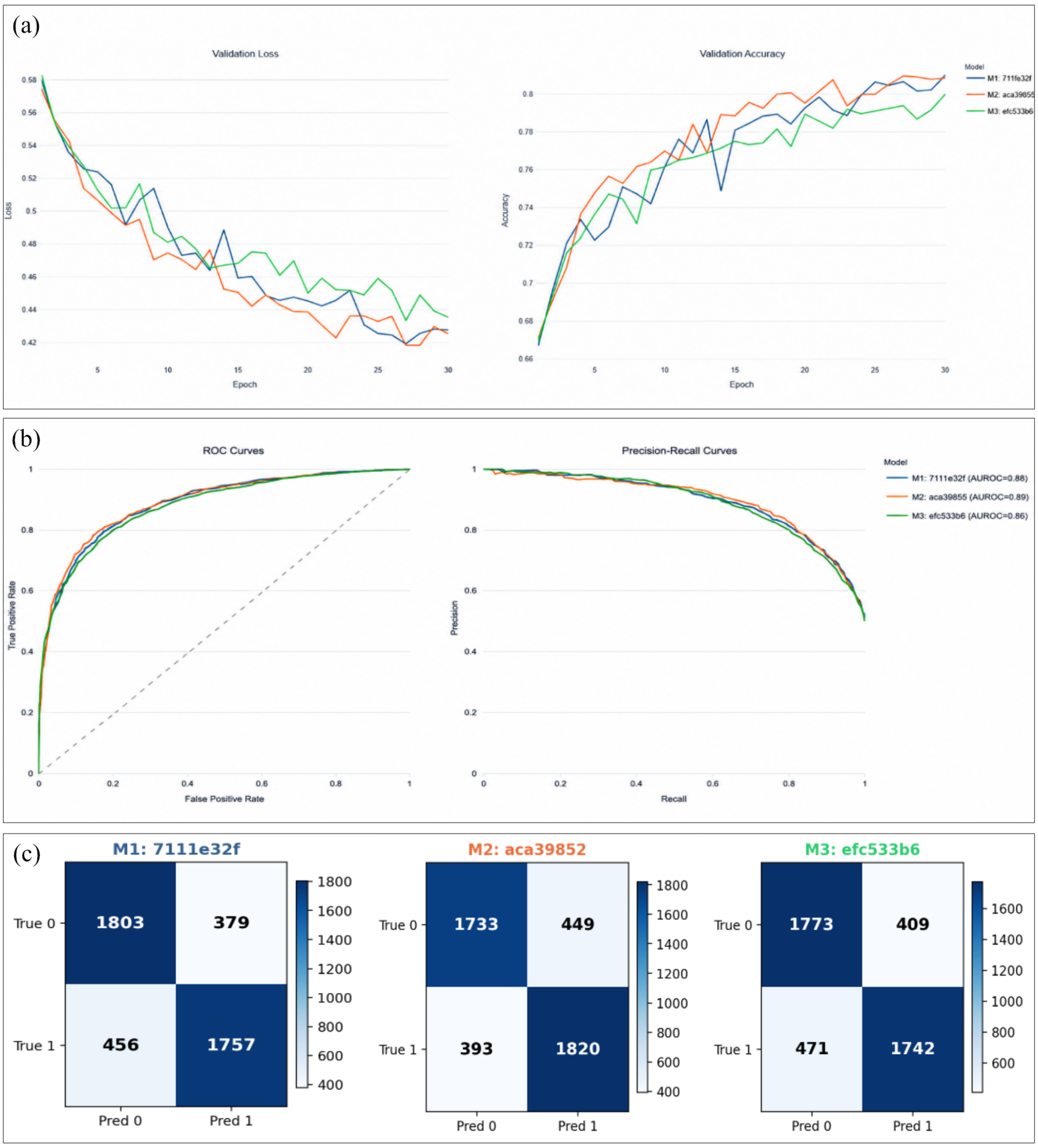

Figure S3: (a)Validation loss (left) and validation accuracy (right) curves across 30 training epochs for M1 (blue), M2 (orange), and M3 (green); (b) ROC curves (left) and Precision-Recall curves (right) for M1, M2, and M3 computed on the validation set. (c) Confusion metrices for M1 (7111e32f, left), M2 (aca39852, center), and M3 (efc533b6, right).

a simpler architecture than M2. The close performance range across all three models (AUROC: 0.8804-0.8893; AUPRC: 0.8924-0.8969) demonstrates that the shared encoder representations are highly informative, and that architectural variation in the fusion head produces moderate but interpretable differences.

Table S4: Validation set performance metrics for M1, M2, and M3 after 30 epochs of training. Highest values per metric are highlighted in green.

| **Metric** | **M1 (7111e32f)** | **M2 (aca39852)** | **M3 (efc533b6)** | **Best Model** |
|---|---|---|---|---|
| Val. Accuracy | 0.8100 | 0.8084 | 0.7998 | M1 |
| AUROC | 0.8874 | 0.8893 | 0.8804 | M2 |
| Avg. Precision (AUPRC) | 0.8957 | 0.8969 | 0.8924 | M2 |
| Precision | 0.8226 | 0.8021 | 0.8099 | M1 |
| Recall | 0.7939 | 0.8224 | 0.7872 | M2 |
| F1 Score | 0.8080 | 0.8121 | 0.7984 | M2 |

##### S2.1.4 Predicted Probability Distribution Analysis

To characterize model calibration and confidence behavior, the platform overlays the predicted probability histograms for all trained models on a shared axis (Figure S4). This overlay is generated automatically upon completion of training and provides an immediate visual summary of decision boundary confidence across architectures. All three models exhibit a pronounced bimodal distribution with substantial probability mass concentrated near 0 and near 1, indicating high classification confidence for the majority of samples. The intermediate probability region (0.2-0.8) is sparsely populated across all models, suggesting minimal ambiguity in the learned decision boundary. Minor distributional differences between models in this intermediate region reflect architectural influences on uncertainty estimation, with M2 displaying a slightly elevated density near $p = 1.0$ consistent with its higher recall. These visualizations are rendered interactively within the platform's Probability Distribution Viewer, accessible without any post-training scripting.

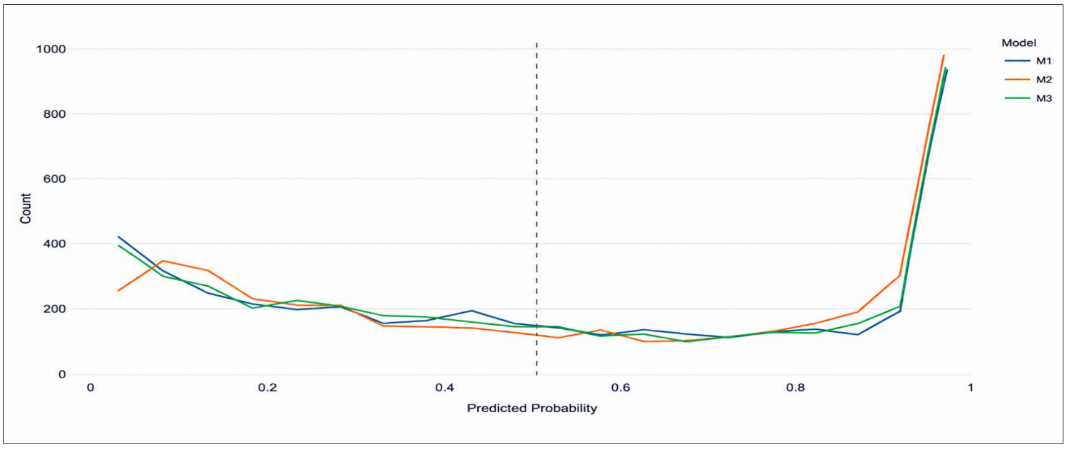

Figure S4: Overlaid predicted probability distribution histograms for M1, M2, and M3. All three models exhibit a characteristic bimodal distribution with pronounced mass near 0 and 1, indicating confident binary discrimination. The dashed vertical line at p = 0.50 marks the classification threshold. M2 displays a marginally higher peak near p 1.0, consistent with its elevated recall.

#### S2.2 Integrative multi-model Inference Analysis

Following model training and registration, the platform's Multi-Model Inference Engine was employed to perform simultaneous inference across three independently trained runs of the same M3 architecture (efc533b6) on a common test dataset. This capability is central to the platform's value proposition: enabling users to assess prediction consistency, identify stochastic variability between training runs, and rank model instances based on aggregate performance, all within a unified graphical interface requiring no external scripting.

##### S2.2.1 Run-Level Classification Performance

Three inference runs (Run 1: 0b1eef2f, Run 2: 897e76c9, Run 3: eddae634) were submitted simultaneously to the inference engine. The classification performance of each run across five metrics computed on the shared test partition is given in Table S5 and presented in Figure S5A. The platform automatically annotates the best-performing run per metric in the comparison table (Table S5) Run 3 (eddae634) achieved the highest scores on all five evaluated metrics: accuracy (0.8123), AUROC (0.8848), AUPRC (0.8994), F1 score (0.8003), and Matthews Correlation Coefficient (MCC = 0.6307). The MCC, which provides a balanced measure of binary classification quality regardless of class size, revealed a meaningful performance advantage for Run 3 (delta = +0.033 relative to Run 2). Run 2 achieved the lowest performance across all metrics, with accuracy 0.7936 and MCC 0.5983, suggesting greater sensitivity to stochastic initialization effects. The platform's automated ranking interface presents these comparisons in tabular form with color-coded annotations, enabling rapid identification of the optimal inference run without manual computation.

Table S5: Multi-run inference performance metrics on the shared test dataset. Run 3 (eddae634) achieved the highest score on all five metrics, as automatically identified and annotated by the Deep-Interact Studio comparison engine.

| **Metric** | **Run 1 (0b1eef2f)** | **Run 2 (897e76c9)** | **Run 3 (eddae634)** |
|---|---|---|---|
| Accuracy | 0.7977 | 0.7936 | **0.8123** |
| AUROC | 0.8768 | 0.8699 | **0.8848** |
| AUPRC | 0.8912 | 0.8887 | **0.8994** |
| F1 Score | 0.7913 | 0.7739 | **0.8003** |
| MCC | 0.5974 | 0.5983 | **0.6307** |

##### S2.2.2 Multi-Run ROC and Precision-Recall Analysis

The platform automatically generates overlaid ROC and Precision-Recall curves for all submitted inference runs, annotated with their respective AUROC values in the legend. Figure S5b presents these curves for the three runs. The ROC curves demonstrate consistent discriminative performance across all three runs, with AUROC values spanning 0.870-0.885. The Precision-Recall curves exhibit greater inter-run divergence at high recall values, consistent with the MCC and F1 differences reported in Table S5. Run 3 maintains superior precision at equivalent recall levels, indicating more effective trade-off management across the decision threshold range. The platform's ability to render these curves simultaneously for multiple runs constitutes a key usability advantage over single-model evaluation pipelines.

##### S2.2.3 Per-Run Confusion Matrices

The confusion matrices for each of the three inference runs are presented in Figure S5C. These matrices are rendered automatically within the platform's result panel upon completion of inference and can be exported individually. Run 3 recorded the lowest false positive count of 331 and the second-lowest false negative count of 700, yielding the most balanced error distribution among the three runs. Run 2 exhibited the highest false negative count of 825 despite achieving a comparatively low false positive rate of 309, reflecting a systematic bias toward the negative class that is directly reflected in its lower recall and F1 score. Run 1 occupied an intermediate position. These distributional differences, while subtle at the metric level, are immediately apparent in the platform's comparative matrix view, illustrating how the interface facilitates rapid, expert-level interpretation of inter-run variability.

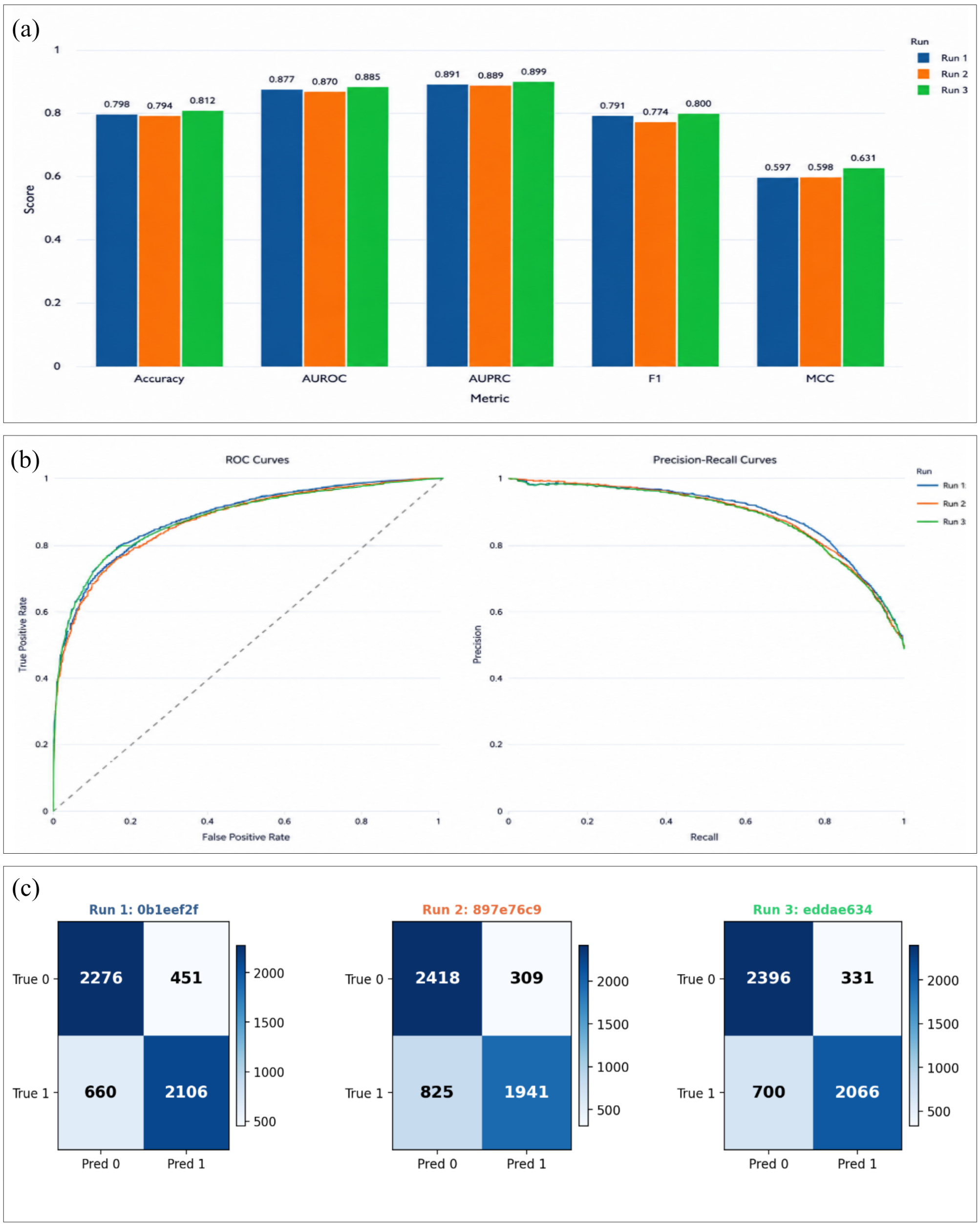

Figure S5: Multi-run inference comparison for three runs on the shared test set, generated automatically by the platform. (a) Grouped bar chart of Accuracy, AUROC, AUPRC, F1, and MCC; Run 3 (green) leads on all five. (b) Overlaid ROC (left) and Precision–Recall (right) curves; Run 3 attains the highest AUROC (0.885) and AUPRC (0.899). (c) Per-run confusion matrices for Run 1, Run 2, and Run 3 (left to right).

##### S2.2.4 Predicted Probability Distribution and Score Scatter Analysis

The platform provides two additional comparative visualizations for multi-run inference, a kernel density estimate (KDE) overlay of predicted probabilities stratified by class label, and a sample-wise probability scatter plot. These tools enable assessment of inter-run calibration consistency and identification of samples with systematically divergent predictions. The KDE plots (Figure S6) reveal that all three runs produce well-separated probability distributions for positive and negative class instances. Positive-class predictions cluster strongly near p = 1.0 for all runs, while negative-class predictions concentrate near p = 0, confirming confident and well-calibrated decision boundaries. Run 3 displays the most pronounced positive-class peak density (<4.0), reflecting higher confidence and tighter probability assignment relative to Runs 1 and 2. Notably, the negative-class KDE curves (dashed lines) are nearly coincident across all three runs, suggesting that inter-run variability is predominantly expressed in positive-class probability estimation rather than negative-class discrimination. The sample-wise scatter plot (Figure S7) reveals pronounced inter-run disagreement for a substantial subset of test samples, particularly in the intermediate probability range ($0.2 < p < 0.8$). While many samples receive consistently high ($p > 0.8$) or low ($p < 0.2$) probability scores across all three runs, a meaningful fraction of pair indices exhibit divergent predictions where one run assigns a high probability while another assigns a low probability. These disagreement instances are quantified by the platform's per-pair disagreement score (standard deviation across runs), the top-ranked values of which are exported to a downloadable CSV (Table S6). This sample-level analysis constitutes a novel capability unavailable in conventional single-model evaluation workflows and provides directly actionable insights for prioritizing experimental validation of uncertain predictions. The top-ranked disagreement sample (sigma = 0.405) illustrates the most informative class of uncertain prediction: Run 3 assigns a high interaction probability (p = 0.948) while Runs 1 and 2 both predict non-interaction (p = 0.412 and p = 0.155, respectively), resulting in a split binary outcome (1 vs. 0 vs. 0). Samples 2–4 exhibit a consistent two-vs-one disagreement pattern, with Runs 1 and 3 predicting interactions while Run 2 assigns low probabilities, suggesting that Run 2's initialization produces a systematically more conservative model. The platform exports this table directly as a CSV file, enabling immediate integration with downstream experimental prioritization pipelines.

Table S6: Top-10 drug–protein pairs ranked by inter-run disagreement score (standard deviation of predicted probabilities across the three inference runs). Pairs with $\sigma > 0.40$ represent high-uncertainty predictions where model runs disagree on classification outcome.

| Rank | Prob (R1) | Pred (R1) | Prob (R2) | Pred (R2) | Prob (R3) | Pred (R3) | Disagreement ($\sigma$) |
|---|---|---|---|---|---|---|---|
| 1 | 0.4115 | 0 | 0.1546 | 0 | 0.9476 | 1 | **0.4046** |
| 2 | 0.9966 | 1 | 0.2983 | 0 | 0.9359 | 1 | **0.3868** |
| 3 | 0.9007 | 1 | 0.1801 | 0 | 0.7743 | 1 | **0.3848** |
| 4 | 0.8162 | 1 | 0.0937 | 0 | 0.6736 | 1 | **0.3827** |
| 5 | 0.9122 | 1 | 0.2718 | 0 | 0.2349 | 0 | **0.3808** |
| 6 | 0.8870 | 1 | 0.2510 | 0 | 0.8883 | 1 | **0.3676** |
| 7 | 0.0943 | 0 | 0.2784 | 0 | 0.8009 | 1 | **0.3666** |
| 8 | 0.1295 | 0 | 0.8036 | 1 | 0.6852 | 1 | **0.3599** |
| 9 | 0.9680 | 1 | 0.3370 | 0 | 0.9476 | 1 | **0.3586** |
| 10 | 0.8128 | 1 | 0.2783 | 0 | 0.1501 | 0 | **0.3515** |

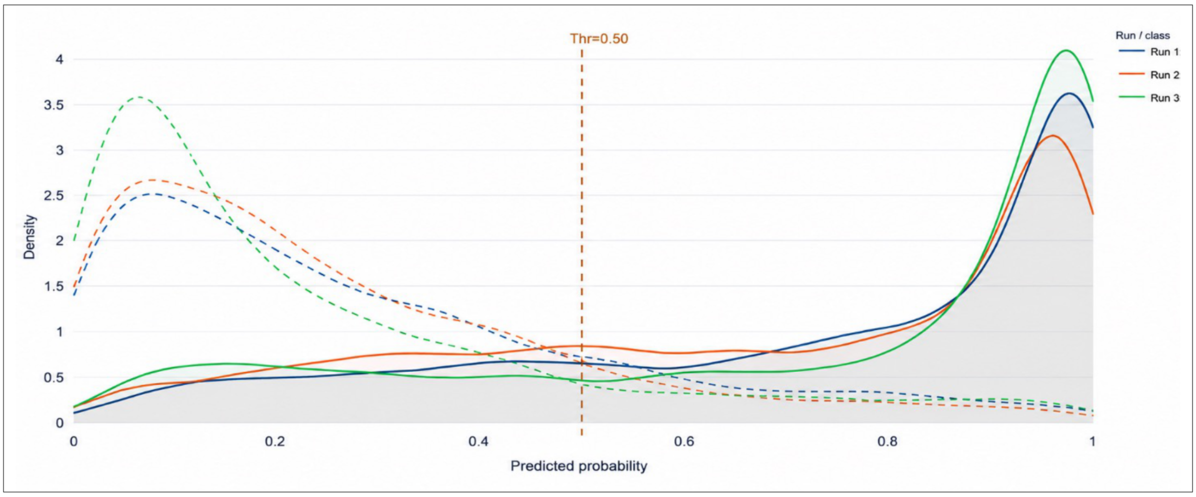

Figure S6: Kernel density estimate (KDE) of predicted probabilities for the three inference runs, stratified by class label (positive class: solid lines; negative class: dashed lines). The vertical dashed line at p = 0.50 marks the classification threshold. All three runs exhibit clear class separation, with positive-class densities peaking near p = 1.0 and negative-class densities concentrated near p = 0. Run 3 (green) displays the sharpest positive-class peak, consistent with its superior calibration and highest AUPRC.

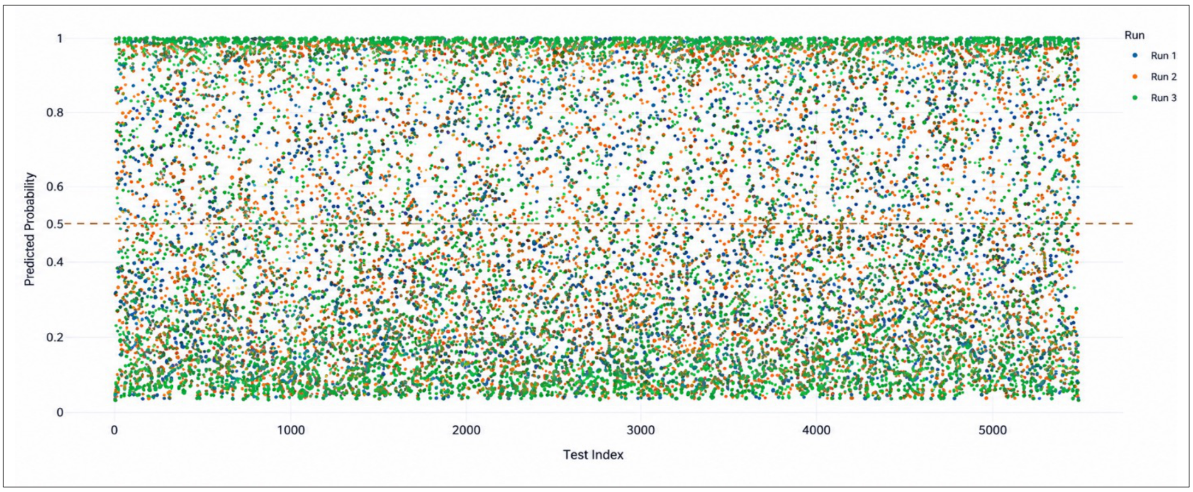

Figure S7: Sample-wise predicted probability scatter plot for all three inference runs across the shared test dataset (5,493 pairs). Each point represents a single drug–protein pair, colored by run identity. The dashed horizontal line at p = 0.50 marks the decision threshold. Substantial vertical scatter between runs for many individual samples reveals high inter-run disagreement for samples with intermediate predicted probabilities, identifying regions of model uncertainty.

#### S2.3 Platform Usability and Functional Summary

The experiments described above were conducted entirely through the Deep-Interact Studio web interface without requiring any command-line interaction, custom scripting, or manual result aggregation. Table S7 summarizes the key platform capabilities exercised during the DTPI case study, illustrating the breadth of functionality available through the graphical user interface. visualization pipeline eliminated the need for post-hoc figure generation, while the integrated export functionality ensured that all numerical results were immediately available in machine-readable format. These characteristics, combined with the reproducibility guaranteed by the platform's hash-based run identification system, demonstrate that Deep-Interact Studio fulfils the requirements of a transparent, usable, and scientifically rigorous tool for comparative deep learning evaluation in bioinformatics research. SHAP-based interpretability analysis (Figure S8A–C) revealed that predictive signal is distributed across both the ChemBERTa compound and ESM-2 protein encoder subspaces, with compound embedding dimension 15 ranking as the single most influential feature and both encoder groups contributing equally at the aggregate level, confirming that the model exploits complementary chemical and sequence-level representations rather than collapsing onto a single modality. Complementary UMAP projections of the raw input embedding space and the model's learned feature space (Figure S8D–E) revealed that while positive and negative pairs are not linearly separable at the pre-model input level, training induces partial geometric separation in the internal representation, with misclassified instances (false positives and false negatives) concentrating at the boundary interface, directly identifying the representational locus of residual prediction error. The platform's network visualization module further translated the inference outputs into an interaction hub network (Figure S8F), identifying high-degree hub proteins with broad predicted target engagement profiles.

Table S7: Summary of Deep-Interact Studio platform capabilities demonstrated in the DTPI case study, illustrating the functional scope available to end users through the graphical interface.

| **Platform Module** | **Functionality Demonstrated** | **Output Generated** |
|---|---|---|
| Model Builder | Simultaneous registration of multiple architectures (MLP, BiLSTM, Residual) | Trained model files, real-time model architecture overview |
| Training Orchestrator | Parallel multi-model training over 30 epochs | Real-time loss/accuracy curves (Fig. S3A) |
| Evaluation Engine | Automated computation of 6 classification metrics per model | Metric comparison table (Table S4) |
| ROC / PR Module | Multi-model AUROC and AUPRC overlay generation | ROC & PR curves (Fig. S3B) |
| Feature embeddings and results embeddings | UMAP projection of raw input embedding space and model learned feature space, colored by split/label and prediction outcome category respectively | Raw embedding UMAP and model feature space UMAP (Fig. S8D–E) |
| Confusion Matrix Viewer | Per-model matrix rendering and export | Confusion matrices (Fig. S3C) |
| Probability Distribution Viewer | Overlaid probability histograms across models | Histogram overlay (Fig. S4) |
| Multi-Run Inference Engine | Simultaneous inference from registered runs | Per-sample probability scores |
| Metric Comparison Bar Chart | Automated grouped bar chart for run comparison | Bar chart (Fig. S5A) |
| Disagreement Analyzer | Per-sample $\sigma$ computation and ranking | Disagreement CSV (Table S6) |
| KDE Viewer | Class-stratified probability density estimation | KDE overlay (Fig. S6) |
| Scatter Plot Module | Sample-wise probability scatter across runs | Scatter plot (Fig. S7) |
| SHAP interpretation | Global and dimension-level SHAP importance across compound and protein encoder subspaces | SHAP spectrum, top-15 ranking, group importance (Fig. S8A–C) |
| Network visualization module | Predicted interaction hub network construction with degree-centrality node sizing | Drug–target hub network (Fig. S8F) |
| Export Engine | One-click CSV/PNG export for all results | Downloadable files (Tables S4–S6) |

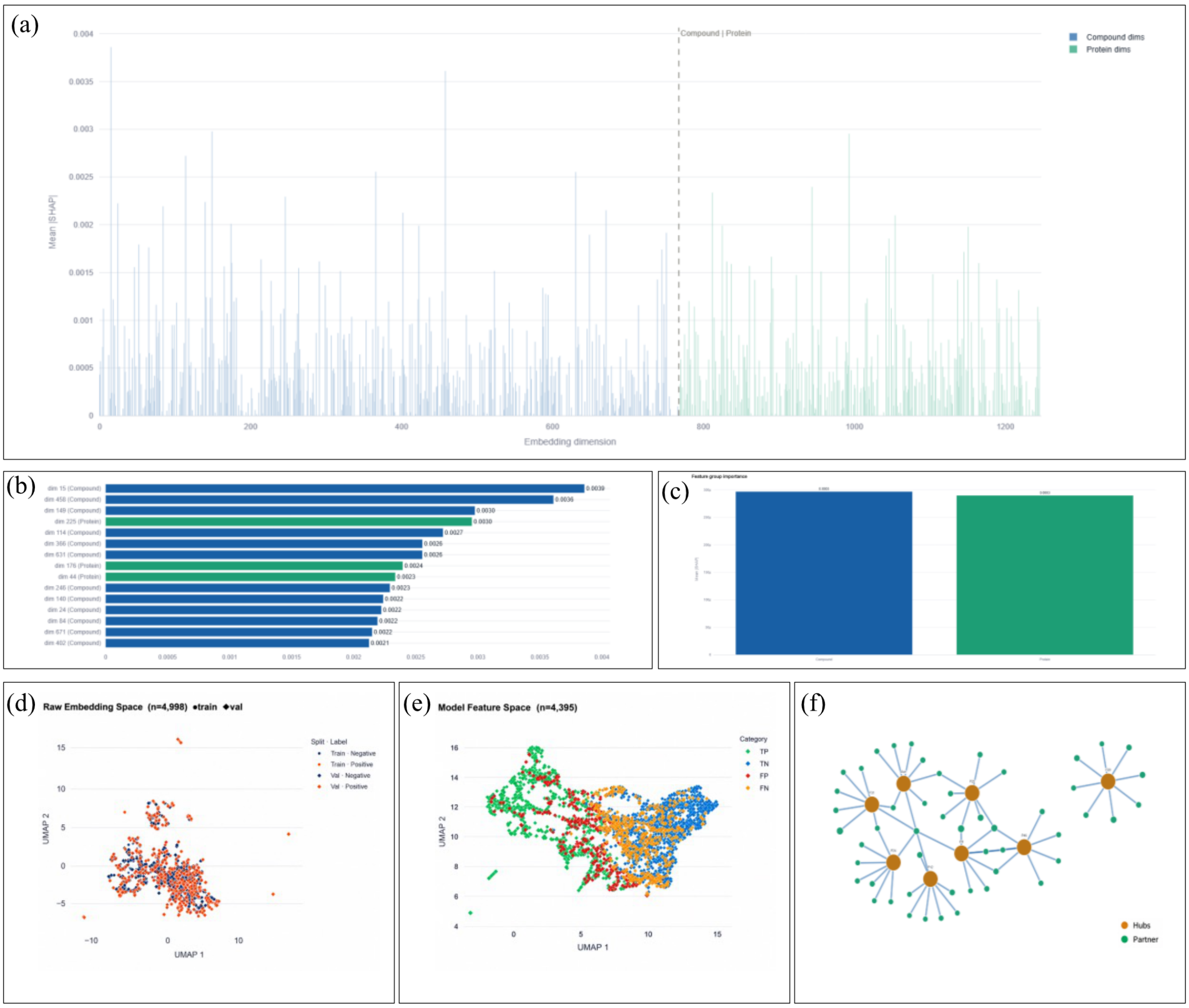

Figure S8: SHAP-based interpretability analysis, embedding space characterization, and predicted interaction hub network generated by the Deep-Interact Studio platform for the DTPI case study. (a) Full-spectrum SHAP importance profile across all embedding dimensions, partitioned into the ChemBERTa compound subspace (blue) and ESM-2 protein subspace (green) (b) Top-15 embedding dimensions ranked by mean SHAP across the test set. (c) Aggregate feature group importance comparing the compound and protein encoder subspaces. Both groups contribute equivalently (d) UMAP projection of the raw input embedding space (train: circles, validation: diamonds; blue = negative, orange = positive). (e) UMAP projection of the model's learned internal feature space, colored by prediction outcome category (TP: green, TN: blue, FP: red, FN: orange) (f) Predicted drug-target interaction hub network for a representative test subset by the platform's network visualization module. Hub protein nodes (orange, scaled by degree centrality) represent targets with the highest predicted interaction counts; compound leaf nodes (green) represent individual drug candidates.

#### S2.4 Comparison with other platforms

The platform extends the capabilities of existing bioinformatics platforms by integrating multi-model training, comparative inference, interpretability, and reproducibility within a unified web-based framework. Unlike current systems that primarily focus on isolated prediction tasks or predefined architectures, Deep-Interact Studio enables user-driven model development across multiple molecular interaction domains, including PPI, DTPI, RPI, and PDI. The platform further incorporates modern representation learning, side-by-side multi-model comparison, and data-centric analytical tools such as feature-space visualization and explainability, thereby supporting transparent, scalable, and reproducible AI-driven bioinformatics research (Table S8).

Table S8: Comparative overview of Deep-Interact Studio and existing bioinformatics platforms highlighting biological scope, multi-model comparative analysis, interpretability, and reproducible AI-driven interaction prediction capabilities.

| **Category** | **Existing Platforms** | **Deep-Interact Studio** |
|---|---|---|
| Accessibility | Primarily web-based platforms focused on predefined workflows with limited user flexibility. | Fully web-based, no-code platform supporting interactive model building, training, inference, and comparison. |
| Biological Scope | Most platforms support only one or two interaction types, mainly PPI or DTPI. | Unified support for multiple biomolecular interaction types, namely PPI, DTPI, RPI, and PDI within a single framework. |
| User-Driven Training | Several platforms provide fixed pre-trained models or limited re-training options. | Supports training on user-uploaded datasets with organism-specific customization and configurable hyperparameters. |
| Representation Learning | Partial support for modern language-model-based encoders such as ProtBERT or ESM. | Integrates state-of-the-art protein and drug encoders including ESM-2 and ChemBERTa. |
| Model Architecture Flexibility | Typically restricted to predefined architectures with minimal customization. | Enables fully configurable layer-wise architecture design and multi-model experimentation. |
| Comparative Analysis | Existing tools mainly evaluate single models independently. | Supports side-by-side multi-model training and sample-level inference comparison across multiple architectures. |
| Interpretability | Limited visualization and explainability support. | Provides dataset statistics, feature-space visualization (UMAP), and explainability-driven analysis. |
| Reproducibility | Reproducibility features are inconsistently implemented across platforms. | Includes centralized model registry, experiment tracking, and exportable reproducible outputs. |
| Scientific Contribution | Focused on isolated prediction or sequence-analysis tasks. | Establishes a unified framework for comparative, interpretable, and reproducible AI-driven interaction analysis in bioinformatics. |

[4] Seyone Chithrananda, Gabriel Grand, and Bharath Ramsundar. Chemberta: large-scale self-supervised pretraining for molecular property prediction. *arXiv preprint arXiv:2010.09885*, 2020.

[5] Xiao-Yong Pan, Ya-Nan Zhang, and Hong-Bin Shen. Large-scale prediction of human protein- protein interactions from amino acid sequence based on latent topic features. *Journal of proteome research*, 9(10):4992–5001, 2010.

[6] Haoquan Liu, Yiren Jian, Chen Zeng, and Yunjie Zhao. Rna-protein interaction prediction using network-guided deep learning. *Communications Biology*, 8(1):247, 2025.

[7] Ningyu Zhong and Zhihua Du. Nasnet-dti: accurate drug–target interaction prediction using heterogeneous graphs and node adaptation. *Briefings in Bioinformatics*, 26(4):bbaf342, 2025.
